# Age-dependent changes in protein incorporation into collagen-rich tissues of mice by in vivo pulsed SILAC labelling

**DOI:** 10.1101/2021.01.13.426496

**Authors:** Yoanna Ariosa-Morejon, Alberto Santos, Roman Fischer, Simon Davis, Philip Charles, Rajesh Thakker, Angus Wann, Tonia L. Vincent

## Abstract

Collagen-rich tissues have poor reparative capacity that is further impaired with age, predisposing to common age-related disorders such as osteoporosis and osteoarthritis. We used in vivo pulsed SILAC labelling to quantify new protein incorporation into cartilage, bone, skin and plasma of mice across the life course. We report highly dynamic matrisome turnover in bone and cartilage during skeletal maturation, which was markedly reduced after skeletal maturity. Comparing young adult with older adult mice, new protein incorporation was reduced in all tissues. STRING clustering revealed epigenetic modulation across all tissues, a decline in chondroprotective growth factors such as FGF2 and TGFb in cartilage, and clusters indicating mitochondrial dysregulation and reduced collagen synthesis in bone. Several of these pathways have been associated with age-related disease. Fewer changes were observed for skin and plasma. This methodology provides dynamic protein data at a tissue level, uncovering age-related molecular changes that may predispose to disease.

## Introduction

As life expectancy extends, the societal burden of age-related diseases is predicted to increase substantially. Ageing, the natural decline of cellular and physiological processes during life, is particularly apparent in collagen rich tissues such as the articular cartilage, bone and skin, leading to osteoarthritis (OA), osteoporosis and impaired cutaneous wound healing in the elderly. Each of these tissues is characterised by an abundance of extracellular matrix (ECM); a dynamic network composed of collagens, proteoglycans, glycoproteins and EMC associated proteins, defined collectively as the matrisome (1). The matrisome constitutes the majority of the tissue by volume; up to 95% in the case of articular cartilage (2). Several mechanisms have been proposed that account for changes to the ECM with age, including reduced matrix turnover, modifications by glycation, crosslinking, proteolytic degradation and accumulation of protein aggregates (3). Such changes affect the regenerative capacity and biophysical properties of the tissues and, in doing so, predispose to disease (4).

Collagens comprise one third of all proteins in mammals and are the major component of connective tissues such as skin, cartilage, bone and tendon. Long lived fibrillar collagens contribute to the accumulation of age-related modifications such as advanced glycation end products (AGEs), protein deamidation or racemization of amino acid residues (5, 6). The long half-life of type II collagen within cartilage is thought to account for the observed limited repair capacity of the tissue (7). In bone, the accumulation of AGEs, combined with collagen loss and cellular senescence, are regarded as critical drivers of osteoporosis (8, 9). Collagens also play an essential role in wound healing; their deposition and quality define wound closure and resulting tensile strength of the scar. Other ECM components have also been implicated in age-related diseases such as cancer and fibrosis (10). Signals from ECM protein fragments and growth factors associated with the ECM (11) can signal through cell-surface receptors to modulate cell proliferation and survival, cell morphology, and tissue metabolism (12).

Although we recognise that ECM remodeling is a unique and important feature of tissues (3), there is still much to understand about the spatio-temporal changes that occur during healthy aging and disease. Most omics studies provide valuable information regarding molecular composition and abundance, but they only represent snapshots of a physiological state at a given time and do not accurately account for dynamic protein turnover within tissues (13). Using radiolabelling methods or by measuring post translational modifications, previous studies in human skin, cartilage, dentin and tendon, have identified major fibrillar collagens as very stable proteins (5, 7, 14, 15), with little measurable incorporation after skeletal maturity (7) and half-lives as long as 117 years in cartilage (type II collagen) and 15 years in skin (type I collagen) (5). Some of these measurements have been based on accumulation of D-aspartate (D-Asp) or advanced glycation end products (AGEs) (5, 15, 16). A limitation of these methods is that time is not the only factor influencing these changes, e.g. high temperatures, common at injury sites, can accelerate racemization (17-19). Metabolic labeling is a more direct method to estimate protein lifespans. Turnover of the major fibrillar collagens in dentin, tendon and articular cartilage is demonstrably very low when measured by the incorporation of ^14^C using the bomb-pulse method; a natural experiment taking advantage of an increase in atmospheric ^14^C after nuclear testing in the 1950s (14, 20, 21). This is in contrast to proteoglycans, which have a high turnover rate (7). This method is suitable for estimating turnover of bulk tissue proteins or single protein types, but is incompatible with mass spectrometry-based proteomics.

Stable isotope labelling (Lys (6)-SILAC-Mouse Diet, SILANTES GmbH), incorporated into the diet overcomes many of the aforementioned problems and has been used to determine incorporation rates of proteins from blood cells, organelles and organs in vivo (22). In this diet, the six naturally abundant ^12^C molecules in Lysine have been replaced with ^13^C, conferring a molecular mass shift of 6 Daltons for each lysine in a proteolytically cleaved peptide. This method was originally used to quantitatively compare the proteome of fully labelled (F2 generation) knockout mice and determine protein dynamics in vivo (22). Quantification of the ‘heavy’ and ‘light’ protein fractions by mass spectrometry over time provides a measure of the incorporation rates of individual proteins (23, 24). In our study we pulse labeled mice with the SILAC diet for three weeks at three different post-natal periods: weeks 4-7 (maximum skeletal growth), weeks 12-15 (young adult) and weeks 42-45 (late adult), then applied mass spectrometry to quantify new protein incorporation into three collagen-rich tissues (articular cartilage, bone and skin) and plasma, allowing us to estimate and compare new protein incorporation rates at proteome-wide scale across the lifecourse.

## Results

### Proteome turnover in cartilage, skin, bone and plasma during skeletal growth

To measure new protein synthesis and incorporation during the period of maximal skeletal growth, we pulse labeled 2 groups mice (groups A and B, Figure 1) by feeding them heavy SILAC diet for 3 weeks from 4 weeks of age. Group A was culled for tissue collection immediately after the feeding period, whilst group B was fed with the light SILAC diet for another 3 weeks (to week 10). Plasma, knee articular cartilage, ventral skin and tibial bone were collected for analysis at week 7 (group A) or week 10 (group B). Tissues were extracted and analyzed by mass spectrometry (Fig. 1).

**Fig 1.**
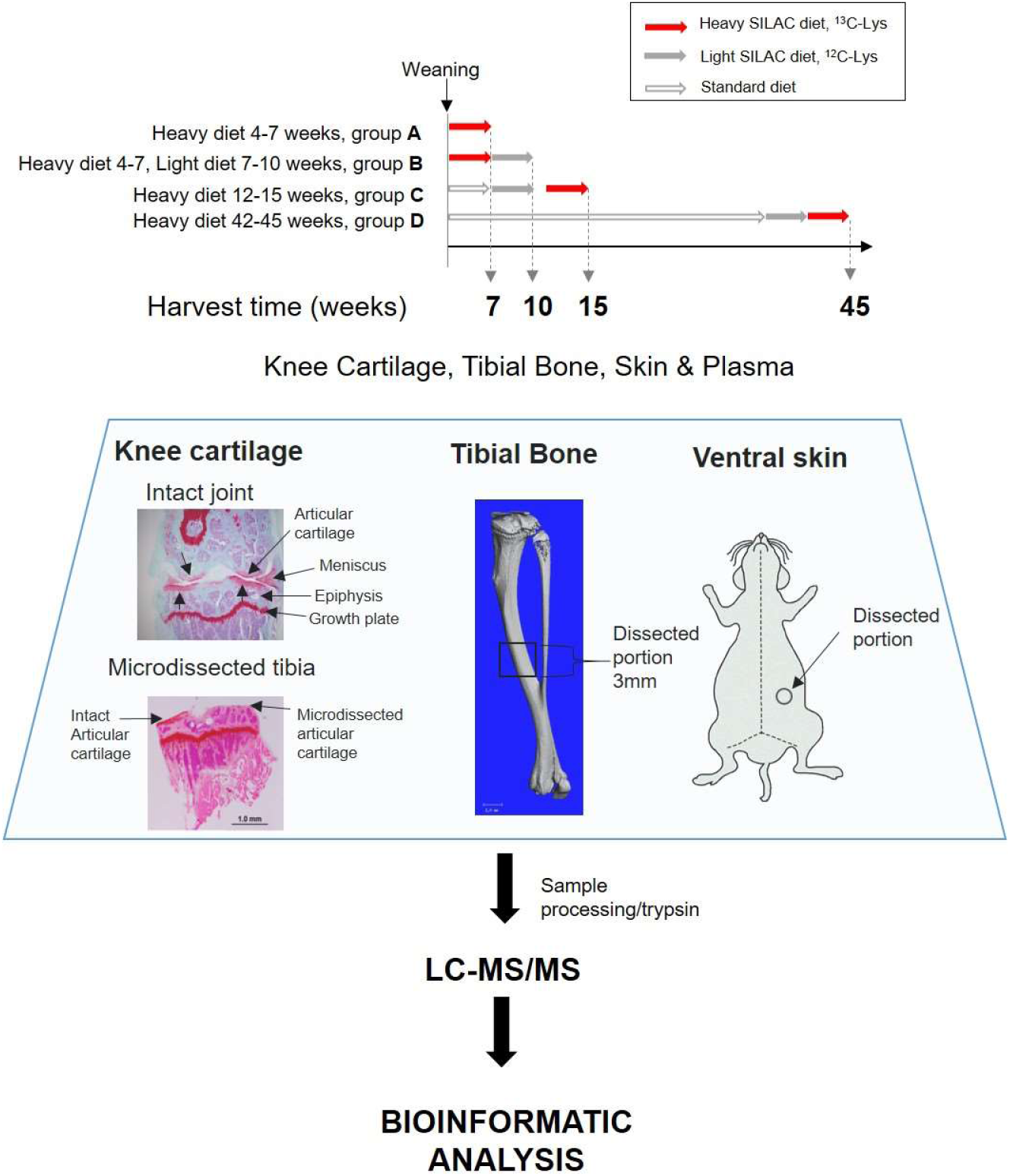
Experimental design. Four groups of four C57BL6 male mice were fed with a heavy SILAC diet (^13^C_6_-Lys) or light SILAC diet (^12^C_6_-Lys) for three weeks at different ages. (A)Two groups of mice (A & B) were fed with the heavy diet from week 4 to 7 of age. Then, group A was culled for tissue collection and B was changed to the light diet from week 7 to 10, then culled for tissue collection. Groups C (week 12 to15 of age) and D (week 42 to 45 of age) were fed with the heavy SILAC diet only. Plasma, knee articular cartilage, trabecular tibial bone and ventral skin were collected from the four mice at the anatomical positions shown above. First panel shows Safranin O stain of intact joint (Upper), and after half right micro-dissection of the articular cartilage (Lower). Tissues were processed, trypsinised and peptides analysed via LC-MS/MS. Peptides and protein identification and heavy/light (H/L) ratios were obtained by Maxquant software.

Fig. 2 shows the percentage of heavy isotope incorporated into individual proteins at the end of the labelling period (week 7) compared with the amount of labelled protein remaining in that tissue after a 3-week washout period of light isotope diet (week 10). In essence, reflecting proteins that are newly incorporated into the tissue and their stability over time. Each tissue was considered separately. We used the plasma profile to define the fast turnover protein group as previously described (25). Proteins that showed over 80% incorporation during the heavy diet period and retained less than 20% after the light diet period, were considered fast turnover proteins (hashed blue area, Fig. 2), while proteins that showed little change in the percentage of isotope incorporation between the end of the heavy diet (week 7) and the end of the light diet (week 10) were regarded as “stable proteins”, (hashed pink area, Fig 2). Over 83% (142/171) of plasma proteins fell within the fast turnover group, while only 3.5% (6/171) proteins were within the stable group. Cartilage showed the lowest proportion of fast turnover proteins (18%, 115/634) when compared with bone (48%, 338/712), skin (39%, 136/352), and was also the tissue that contained more proteins with less than 40% of isotope incorporation during the heavy diet period (bottom left hand corner of graphs, Fig. 2). The patterns of heavy isotope incorporation were similar across the 4 biological replicates for each tissue and time point, (Fig. S1). The full profile of the 4 tissue proteomes during skeletal growth are presented in Table S1.

**Fig 2.**
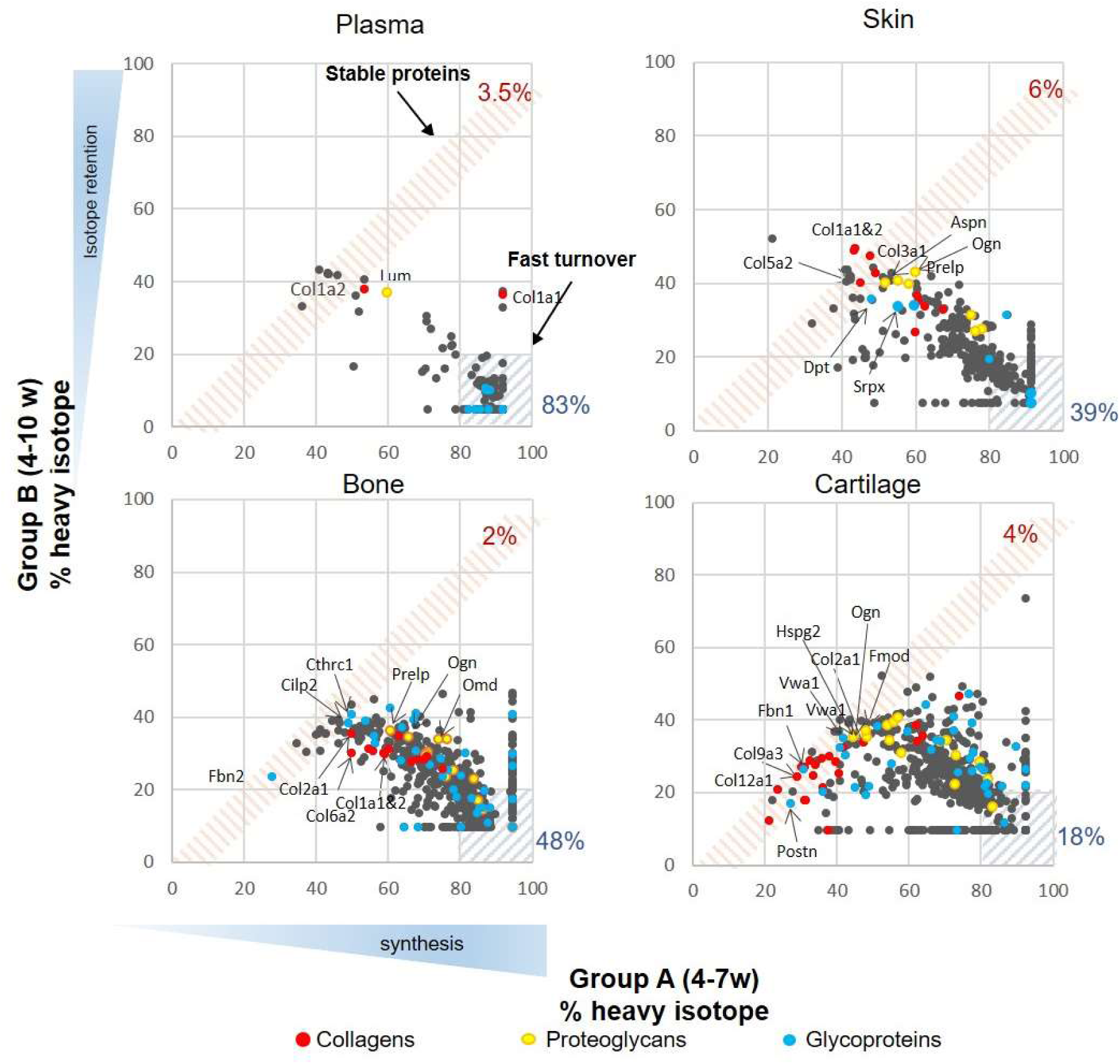
Synthesis rate and incorporation in plasma, skin, bone and articular cartilage during skeletal growth. Each dot represents the mean of the percentage of incorporation of the heavy isotope for an individual protein, n=4. The x axis represents percentage of the heavy isotope incorporation into proteins from week 4-7 of age (group A) and the y axis the heavy isotope subsequent loss during the light diet, week 7-10 of age (group B). Collagens are highlighted in red, proteoglycans in yellow and glycoproteins in blue. Percentage of stable and fast turnover proteins are stated next to hashed squares and parallelograms. Stable (S): plasma 3.5% (6/151), fast turnover (FT) 83% (142/171), S in skin: 6% (20/352), FT: 39% (136/352); S in bone: 2% (16/712), FT: 48% (338/712), S in cartilage: 4% (28/634), FT: 18% (115/634).

Regarding matrisome proteins, the three most stable collagens (red dot), proteoglycans (yellow dot) and glycoproteins (blue dots) are highlighted for each tissue (Fig. 2). During this period of skeletal growth, collagens exhibited variable stability across all tissues, and this included the fibrillar collagens (Type I, II, III, V and XI), which are generally regarded as being the most stable. Proteoglycan synthesis was more dynamic, showing less isotope retention after the washout period (smallest p values, unpaired t-tests). Glycoproteins spanned a wide turnover range in all tissues.

### Age-dependent remodelling of the tissue proteome

To investigate how new protein synthesis and incorporation changes during ageing, we pulse labeled another 2 groups of four male mice: group C and D. Group C represented skeletally mature (young adult) mice fed heavy diet between 12-15 weeks of age and group D mice fed heavy diet between 42-45 weeks of age (late adult). For each animal, plasma, knee articular cartilage, ventral skin and tibial bone were processed immediately after the 3-week heavy diet as in A (Fig. 1). Histograms showing the frequency of incorporation of the heavy isotope into proteins in groups A, C and D are shown in Fig. S2. Protein synthesis generally decreased in all 4 tissues with age, as shown by a shift to the left of the histogram bins. As expected, for bone and cartilage new protein incorporation decreased more between skeletally immature (Group A) and young adult animals (Group C) than between young (Group C) and late adult mice (Group D), commensurate with cessation of skeletal maturation. Although all tissues showed an age-related decline in new protein incorporation, protein synthesis was significantly different between the four tissues and at each age group, Kruskal-Wallis test, p<0.0001 for all comparisons. Cartilage and bone continued to see a decline between the two adult groups, while skin and plasma maintained a higher basal synthesis level that was only minimally affected over this 30 week period.

We were able to determine differences in protein synthesis rates in 27 different collagens α chains (Fig. 3, Table S2), 21 different proteoglycans (Fig. 4, Table S3) and 66 glycoproteins (Fig S3, Table S4) across all tissues and age groups. Fibrillar collagens showed the greatest decline of incorporation rates between skeletal immaturity and maturity, compared with non-fibrillar: FACIT (fibrillar associated with interrupted triple helices) (IX, XII, XIV, XVI, XXII), network (IV, X) and beaded filament (VI) collagens. Major collagens, type II in cartilage and type I in bone and skin, had the most dramatic decrease in new incorporation, falling to <2% in cartilage upon reaching skeletal maturity. Compared with collagens, proteoglycans and glycoproteins maintained a higher incorporation rate across the life course.

**Fig 3.**
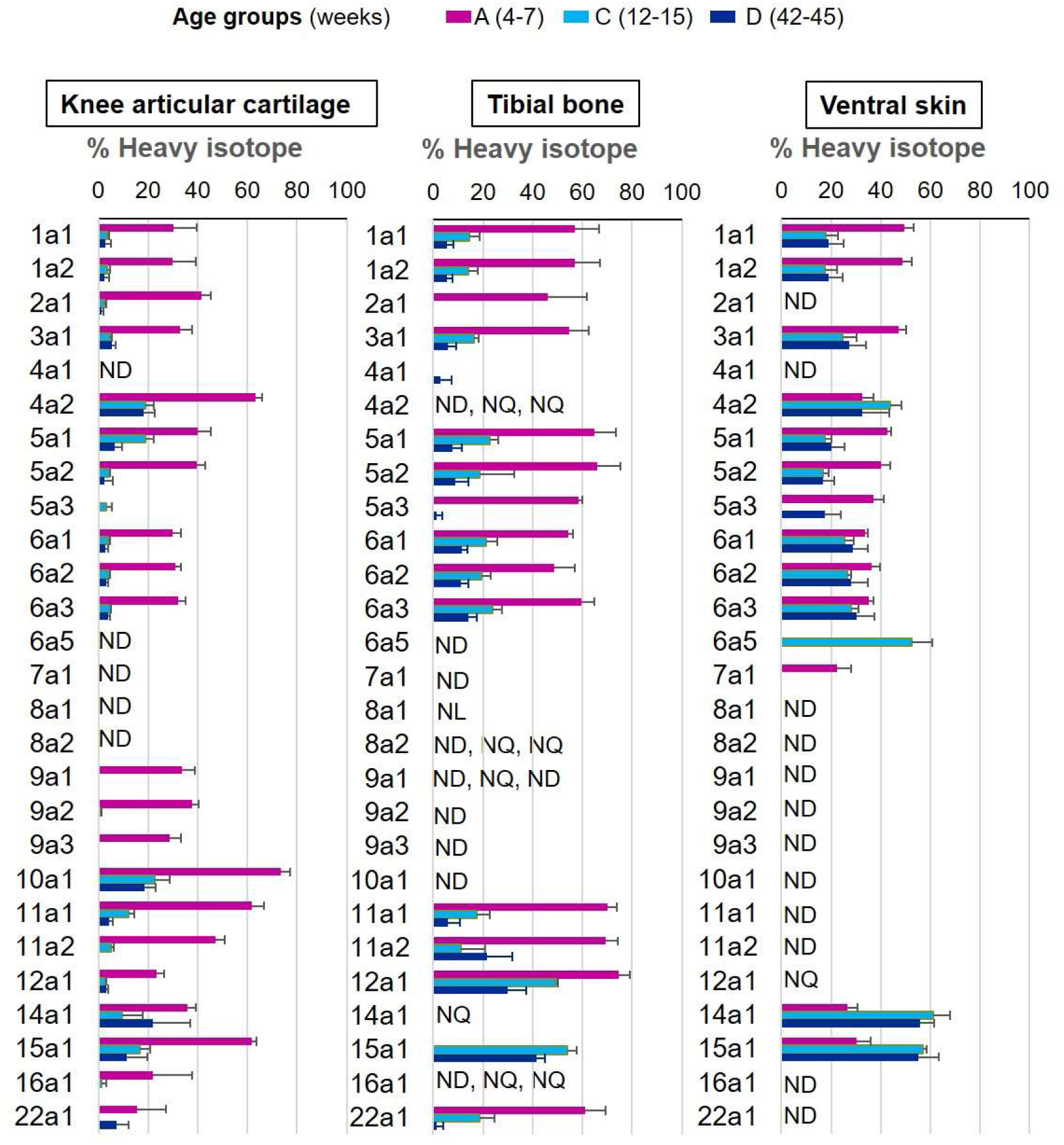
New collagen incorporation into different tissues during aging. The percentage of newly synthesized collagen incorporated into articular cartilage, tibial bone and ventral skin is estimated by the incorporation of the heavy isotope (^13^C_6_-Lys) into proteins during the three weeks of heavy diet. The three periods of heavy diet were compared: skeletal growth (4-7 weeks old), young adults just after skeletal maturity (12-15 weeks old) and older adults (42-45 weeks old). Error bars represent the standard deviation (n=4). ND= not detected, protein group not present in the group dataset. NQ= not quantified, protein group present, but with <2 samples quantified. NL= not labeled, protein group with only light isotope quantified.

**Fig 4.**
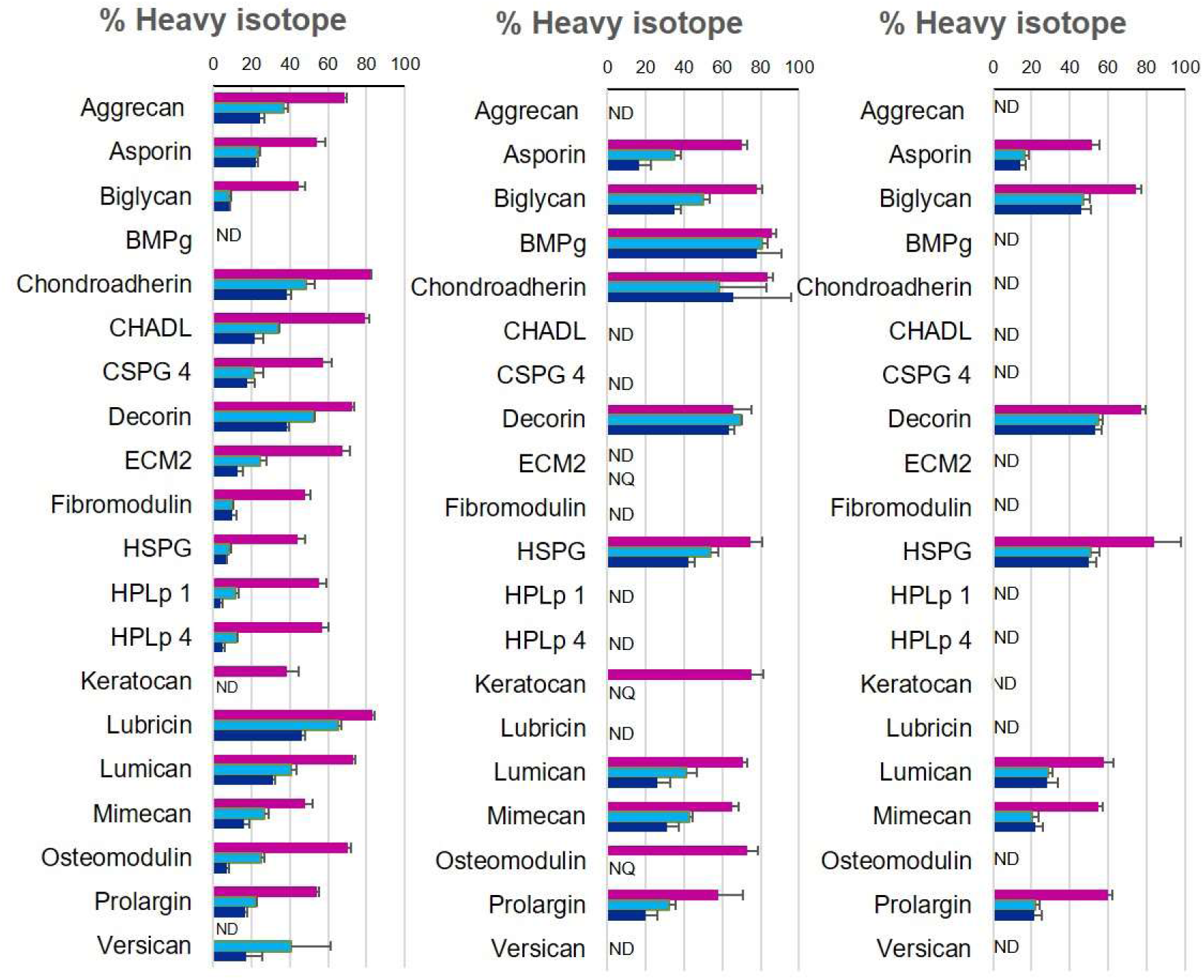
New proteoglycan incorporation rate into different tissues during aging. The percentage of newly synthesized proteoglycans incorporated into articular cartilage, tibial bone and ventral skin is estimated by the percentage of incorporation of the heavy isotope (^13^C_6_-Lys) into the proteins during the three weeks of heavy diet. Protein synthesis and incorporation was estimated across life, covering skeletal growth (4-7 weeks old), young adults just after skeletal maturity (12-15 weeks old) and older adults (42-45 weeks old). Error bars represent the standard deviation (n=4). ND= not detected, protein group not present in the group dataset. NQ= not quantified, protein group present, but with <2 quantified samples. BMPg, bone marrow proteoglycan.

To examine changes in new protein incorporation across the whole tissue proteome with age, we compared the whole proteome profile in skeletally mature groups D and C. Data are represented in volcano plots (Fig. 5 A-D). Significantly regulated proteins with –1.5 >FC> 1.5, BH correction FDR<0.05, for each tissue are highlighted (bold points). Overall, the proportion of newly synthesised proteins within each tissue decreased with age, and only a few proteins had increased incorporation with age in all four tissues. In plasma, 83 out of 180 proteins exhibited statistical significant regulation upon ageing. In connective tissues these were 37 from 452 in skin, 291 from 572 in bone, and 175 from 597 in cartilage (Proteins shown in Table S5A-D). Only 2 regulated proteins were common among the three collagenous tissues; H1f0, a histone protein, downregulated in cartilage and bone but upregulated in skin, and Rpl12, a structural constituent of the ribosome 60 S subunit, downregulated in the three tissues. Articular cartilage and trabecular bone proteomes shared 9.6% of the statistical significantly regulated proteins while overlap between bone and skin was 1.1%, and cartilage and skin 0.7%.

**Fig 5.**
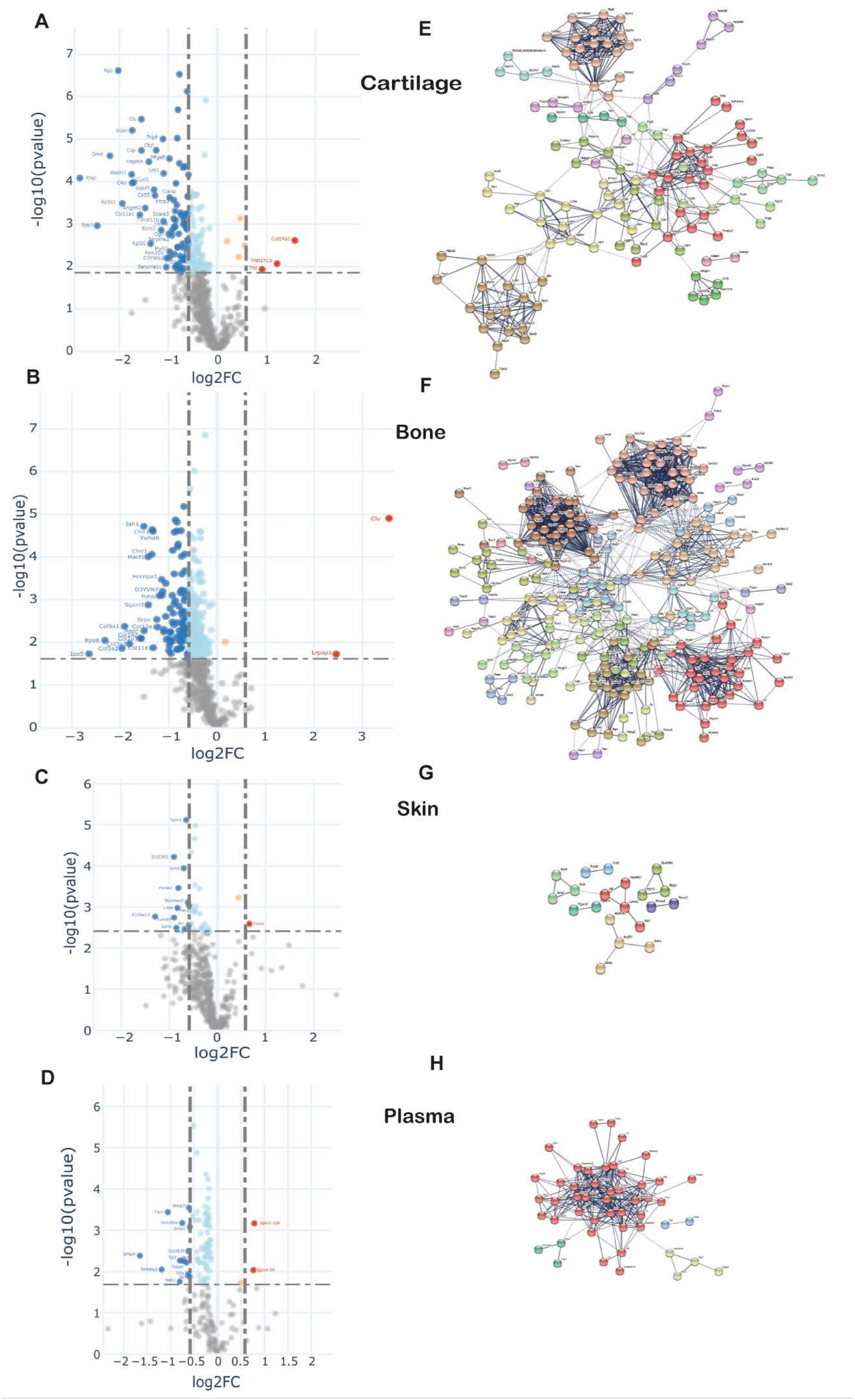
Changes in protein synthesis and incorporation rates during tissue remodelling. **A-D**. Differential protein incorporation rates in young versus older adult tissues. A-cartilage, B-bone, C-skin and D-plasma. Volcano plots, unpaired ttest with BH correction FDR<0.05, FC>1.5, n=4. Full list of proteins available in table S5. **E-H**. STRING cluster analysis of differentially incorporated proteins in each tissue. E-cartilage, F-bone, G-skin, F-plasma. Edges show high confidence interaction score=0.7. Networks clustered to MCL inflation parameter=2. Cluster elements are listed in table S6.

STRING protein interaction networks of statistical significantly regulated proteins for the four tissues are shown in Fig. 5 E-H, with full list of cluster elements in Table S6A-D. Each of the tissues shared a common cluster relating to regulatory elements involving ribosomal and histone proteins, albeit with different constituent proteins in each tissue. The largest cluster of interaction networks in cartilage (Fig 5E) was mainly formed by extracellular matrix proteins such as collagens, collagen processing-related and proteoglycans. Also notable, were 2 clusters abundant in proteoglycans and growth factors, and another cluster rich in myosin family members. 15 proteins with decreased incorporation have previously been implicated in osteoarthritis development (Table S5A).

In bone, the largest cluster represented several myosin family members, involved in muscle contraction, some structural proteins and proteins with roles in ATP transfer. Two unique trabecular bone clusters were involved in energy metabolism; one formed by 29 mitochondrial proteins such as ATP synthases, cytochrome oxidases and NADH dehydrogenases, and the other largely by enzymes involved in glycolysis and fatty acid metabolism (Fig. 5F, Table S6B). Three proteins with statistically significant decreased incorporation have previously been linked to osteoporosis, Biglycan, Col1a2 and Troponin C2 (Table S5B), and SNPs in the genes of two further proteins, also with decreased incorporation, YWHAE and CD44, have been found associated with bone mineral density (BMD), increase bone fractures in mouse models (YWHAE) and osteoclast activity (CD44).

One striking result was the relative paucity of the percentage of age-regulated proteins in skin (8%) compared with cartilage (29%) and bone (51%) despite a similar number of total proteins quantified (Fig. S4). In skin a number of weak clusters were identified with a couple involving energy metabolism (Fig 5G, Table S6C). One downregulated protein in the skin, solute carrier family 3 (SLC3A2) (Table S5C), has been implicated in wound healing (26, 27).

In plasma, one strong cluster comprising 40 proteins with roles in lipid transport (apolipoproteins), proteins involved in the innate immune response, and several serpin family members, were identified (Fig 5H, Table S6D).

## Discussion

To extend our knowledge of connective tissue renewal during growth and ageing, we used metabolic labeling with stable isotopes in combination with quantitative proteomics to estimate individual protein turnover and incorporation rates in live animals across the life course. Here, we describe comprehensive tissue proteome incorporation rates for proteins in skin, cartilage, bone and plasma at 3 postnatal stages of life: during maximum skeletal growth, and at young and late adulthood. Our study shows that although new protein incorporation decreased significantly in all tissues after skeletal maturity, it displays distinct temporal and molecular tissue signatures.

The issue turnover can be divided into two stages: tissue modelling during development and growth, and tissue remodelling, where damaged proteins are replaced to maintain homeostasis and functionality (28, 29). Here, we measured turnover during the modelling phase using a SILAC pulse-chase strategy (groups A and B) and remodelling by measuring SILAC incorporation rates (groups C and D). In general, incorporation rates in all four tissues decreased as animals aged, but the decline was most prominent for cartilage and bone. We found that cartilage and bone had the largest synthetic decline once skeletally mature (young to older adulthood), with fibrillar collagen incorporation rates dropping to <2% in cartilage (type II collagen), and around 5% in bone (type I collagen). These results are in line with previous observations showing minimal turnover of type II collagen after skeletal maturity in human articular cartilage (7), perhaps accounting for the low tissue repair capacity of this tissue. Bone was the only tissue that showed a strong collagen cluster (enriched in 12 collagens) when comparing young with older adult tissue, possibly indicating that bone retains a good synthetic response in young adults and this only becomes significantly blunted later in life. In skin, significant decreases were also observed in fibrillar collagens, but the transition between skeletally immature and mature tissues were less marked, with substantial on-going incorporation of fibrillar collagens through young and older adulthood. This is in line with previous studies that have determined skin to be structurally mature at birth and able to renew throughout the life course (30).

Previous age-related research has identified several cellular pathways that are consistently found to be dysregulated during ageing and in age-related disease. Cell energy and metabolism, including mitochondrial dysfunction, is one of the top processes (31-33). Proteins such as clusterin, superoxide dismutase 1(SOD1), and apolipoprotein E have previously been found to be dysregulated in ageing (34-36). In this study, we found clusterin dysregulation in plasma, cartilage and bone of 45 week old mice when compared with those at 15 weeks of age, and downregulated SOD1 in cartilage. Modulators of protein synthesis such as heterogeneous nuclear ribonucleoprotein D, that affects stability and metabolism of mRNAs, were also down-regulated in cartilage, and have previously been linked to ageing (37). Our data showed only 2 proteins where reduced incorporation was seen in all three collagenous tissues. These included the ribosomal protein, RPL12 and Histone H1.0. However, strong down-regulated clusters, rich in ribosomal proteins were present in bone, cartilage and skin, indicating a decline in incorporation rates of ribosomal structural proteins across all tissues as the animal ages. This finding is in agreement with several studies in which progressive decreases in the expression of ribosomal proteins or rRNA has been observed during ageing (38, 39).

One of the aspects of tissue remodelling is the ability to repair or adapt a damaged tissue to restore homeostasis. Collagen-rich tissues are believed to have a limited reparative capacity, mainly due to the long half-life of fibrillar collagens that are integral to the structure and strength of the tissue (5). Our results identified new incorporation of multiple collagens in each of the connective tissues, with almost all of them showing decreases in incorporation with age, although tissue specific temporal differences were apparent. For instance, bone was the only tissue that showed a strong collagen cluster (enriched in 12 collagens) when comparing young with late adult tissue.. These results might be interpreted as indicating that bone retains good synthetic response in young adult and this only becomes significantly blunted later in life. In cartilage fibrillar collagen synthesis was significantly compromised as soon as skeletal maturity was reached. In skin that synthetic response was maintained for much of adult life as we might expect skin repair responses to be.

Anabolic and repair processes are likely to impact directly on age-related diseases. Interestingly, in cartilage 15 of the downregulated proteins have been associated with osteoarthritis. Several of these have known protective functions in cartilage tissue homeostasis, such as Prg4 (also known as lubricin) (40), Timp3, an inhibitor of disease-associated metalloproteinases (41), and LDL receptor related protein 1(Lrp1), an important chondrocyte scavenger receptor (42). In addition, a number of pro-repair and chondrogenic growth factors were reduced in late adult cartilage such as fibroblast growth factor 2 (FGF2) (43) connective tissue growth factor (CTGF) (44) and transforming growth factor b (TGFb1) (45, 46). Frzb, an antagonist of the Wnt signalling pathway, has also been implicated in osteoarthritis development (47-49). These results not only suggest an explanation for why age predisposes individuals to osteoarthritis development, but also point towards loss of pro-repair pathways in the pathogenesis of age-related OA. This conclusion is supported by recent genome wide association studies (50).

The balance between bone formation and resorption is compromised with increasing age, resulting in progressive net bone loss (51). In trabecular bone we observed decline in several collagens and proteoglycans, which are major structural components of the ECM. Bone also displayed a very strong metabolic, mitochondrial signature that has been linked to many ageing processes (32, 33). Three proteins whose incorporation rates decreased with age are regarded as biomarkers of osteoporosis. One of these was biglycan, a bone modifying protein in murine osteoporosis (52). Collagen 1a2, the most abundant protein in bone matrix, also showed a statistically significant reduction in late adult mouse bone. Degradation fragments of collagen 1 (CTX-1) are regarded as a key biomarker in the diagnosis and treatment of osteoporosis (53). In addition, we found decreased incorporation in two proteins, 14-3-3 protein epsilon and cd44. SNPs in these two protein coding genes, YWHAE and CD44, have been found associated with bone mineral density for the first time in two recent GWAS studies (54) (55).

In skin, only 8% of proteins had altered incorporation rates in the older group, considerably lower than in cartilage and bone (29% and 51% respectively). 6 clusters comprising a maximum of 4 proteins in each were discerned. Wound healing is known to be impaired in the elderly but this phenotype is probably less apparent in ‘middle aged’ individuals which would better represent our older age mice (56). One protein, solute carrier family 3 member 2 (SLC3A2), part of an amino acid transporter, showed reduced protein incorporation in the older group. Deletion of SLC3A2 in murine epidermis impairs epidermal wound healing (57), and has been associated with ageing (58). Although there were several proteins with statistically significant differences in incorporation rates in older skin, the clusters didn’t predict an overall negative effect on wound healing.

We recognise a number of limitations in this study. For cartilage and bone, an insoluble pellet remained after sample processing. It is likely that the insoluble fraction is mainly composed of the oldest, highly cross-linked portion of major fibrillar collagens that increases with age (5, 59), e.g. type II in cartilage and type I (plus mineral components) in bone. In cartilage, the contribution of collagen to the total amount of protein measured by mass spectrometry decreased by 8% between skeletally immature and skeletally mature animals, possibly due to a higher percentage of insoluble cross-linked fibrillar collagen in the older tissue. Therefore, in the older group, the proportion of newly synthesised collagens might be overestimated. This problem was not evident for skin, which solubilised completely during sample processing. Another limitation was that we only studied mice until 45 weeks of age, approximately half the lifetime of a laboratory mouse. We did not include older mice because they frequently develop spontaneous osteoarthritis, which would have been a co-founding variable in cartilage, preventing us from discerning effects due to age and those due to disease. This problem was probably not so relevant in bone, where loss of mineral density may occur insidiously with age but clinically significant disease would be unusual without estrogen deficiency (in females) or immobilisation (60). Finally, we limited this analysis to only three contrasting connective tissues. As the labelled mouse carcasses are currently frozen, the authors welcome collaborative approaches to extend this analysis to other tissues in line with the 3R principles (61).

Compared with protein abundance, that only captures a snapshot of a protein at a particular time, this methodology provides insights into dynamic protein turnover in different tissues and at different ages. In this study, we reveal distinct age-dependent molecular signatures in connective tissues and identify shared and tissue specific protein clusters that could represent novel pathway targets for age-dependent disease.

## Materials and Methods

### Samples and Reagents

Wild-type C57Bl/6 mice, ages 4, 11 and 41 weeks, were obtained either from an in-house colony or from Charles River, Oxford, England. ^13^C_6_-Lysine-SILAC (97 % atom ^13^C)- (heavy diet) and ^12^C_6_-Lysine-SILAC (light diet) were purchased from SILANTES GmbH, Munich, Germany. Animals’ care was in accordance with institutional guidelines.

### Pulsed SILAC Labeling experimental design

Our labeling strategy was aimed at labeling long lived ECM proteins. Therefore, four groups of four male mice were fed with a stable isotope diet for a period of three weeks at different ages that spanned from weaning to late adulthood. The labeling scheme (fig 1A) was as follows: Two groups of mice (A and B) were fed with the heavy diet from week 4 to 7. Then, one of these groups was culled for tissue collection and the other was changed to the light isotope diet (^12^C_6_-Lys) from week 7 to 10, then culled for tissue collection. This was used to calculate proteome turnover rates during skeletal growth. The third group (C) was fed with the heavy isotope diet from week 12 to 15, (young adult), and the fourth group (D) was fed with the heavy isotope diet from week 42 to 45, (late adult). All mice were acclimatised to the SILAC diet formulation by feeding them with the light isotope diet for 1 week before introducing the heavy diet. All mice gained weight as expected (data not shown).

### Tissue harvest

At the end of each labeling period, corresponding to ages 7, 10, 15 and 45 weeks, animals were culled injecting a terminal dose (30mg/animal) of the anaesthetic Pentoject. Blood, knee articular cartilage, tibial trabecular bone and ventral skin were collected from each animal (n=4 mice per time point) as follows: blood was collected by cardiac puncture and mixed with 0.5 M anticoagulant EDTA to obtain a final concentration of 5 mM EDTA. The buffered blood was centrifuged for 15 minutes at 3000 rpm at 4 degrees Celsius. The resulting supernatant (plasma) was transferred to a 0.5 ml tube to be used for analysis. Knee articular cartilage was harvested using a micro-dissection technique previously developed described by our group (62). Articular cartilage from the femoral and tibial surfaces of both knees of one mouse were micro-dissected under the stereo microscope and collected in 1.5 ml micro-centrifuge tubes containing 50 αl PBS. Tibial trabecular bone samples were sectioned between the crest of the tibia and the fibula. The bone marrow was flushed with PBS using a 25 G needle. Approximately 0.8 cm^2^ of skin was cut from the lower flank of the ventral surface. Hair was removed using hair removal cream (Veet sensitive skin). Adipose tissue and blood vessels were removed from the subcutaneous region under the stereo microscope and a portion of clean skin was collected using a 6 mm biopsy punch. All tissues were washed with phosphate buffered saline (PBS [pH 7.4]), and stored frozen at −80 °C until further processing.

### Mass spectrometry sample preparation

Bone and cartilage samples were placed in 180 µL of 5 mM dithiothreitol (DTT) and heated at 65 °C for 15 minutes. After cooling to room temperature, samples were alkylated with 20 mM iodoacetamide (IAA) for 30 minutes. To quench remaining IAA, DTT was added and samples were incubated for 30 minutes. Then, samples were incubated with 4M GuHCl for 2h. The samples were adjusted to pH 8 with 400 mM tris base. Proteins were digested with 1 µg of trypsin overnight at 37 °C. Digestion was terminated adding trifluoroacetic acid (TFA) to a final concentration of 0.5-1 %. Peptides were purified using C18 solid phase extraction cartridges (SOLA HRP SPE cartridges, Thermo Fisher Scientific).

5 µl of plasma was reduced with 5 mM DTT, alkylated with 20 mM IAA and proteins precipitated with methanol/chloroform (63). Precipitated proteins were solubilised in 6 M urea buffer. Urea was diluted to <1 M with milli-Q water and proteins digested with trypsin overnight at 37°C at an enzyme to substrate ratio of 1:25. Digestion was terminated adding TFA to a final concentration of 0.5-1%. Peptides were purified using C18 solid phase extraction cartridges as above.

Skin samples were grinded in a Cryomill, then incubated in 8 M urea, 3 % SDS with protease inhibitors (cOmplete Mini EDTA-free Protease Inhibitor Cocktail, Roche) at room temperature for 1 hour in a total volume of 3 mL. As there was still a substantial pellet, samples were shaken overnight at 4 °C to solubilise the pellet. Samples were centrifuged at 2,500 *g* for 10 minutes and 100 µL of sample was taken for further processing. Proteins were reduced with 5 mM DTT for 30 minutes at room temperature, alkylated with 20 mM IAA for 30 minutes at room temperature and precipitated with methanol/chloroform. Precipitated proteins were solubilised in 6 M urea buffer. Urea was diluted to <1 M with milli-Q water and proteins digested with trypsin overnight at 37°C at an enzyme to substrate ratio of 1:25. Digestion was terminated adding TFA to a final concentration of 0.5-1 %. Peptides were purified using C18 solid phase extraction cartridges as above.

### Liquid Chromatography-Tandem Mass Spectrometry (LC-MS/MS)

Tissues from groups A and B as well as skin samples were analysed on a LC-MS/MS platform consisting of Orbitrap Fusion Lumos coupled to an UPLC ultimate 3000 RSLCnano (Thermo Fisher Scientific) and samples from groups C and D with similar platform but coupled to a Q-Exactive HF. Samples were loaded in 1 % acetonitrile and 0.1 % TFA and eluted with a gradient from 2 to 35 % acetonitrile in 0.1 % formic acid and 5 % DMSO in 60 minutes with a flow rate of 250 nL/min on a 50 cm EASY-Spray column (ES803, Thermo Fisher Scientific).

### Orbitrap Fusion Lumos

The survey scan was acquired at a resolution of 120,000 between 400-1500 m/z and an AGC target of 4E5. Selected precursor ions were isolated in the quadrupole with a mass isolation window of 1.6 Th and analysed after CID fragmentation at 35 % normalized collision energy in the linear ion trap in rapid scan mode. The duty cycle was fixed at 3 seconds with a maximum injection time of 300 ms, AGC target of 3E4 and parallelization enabled. Selected precursor masses were excluded for the following 60 seconds.

### Q-Exactive HF

The survey scan was acquired at a resolution of 60,000 between 375 – 1500 m/z with an AGC target of 3E6, up to the top 12 most abundant ions were selected for fragmentation from each scan. Selected precursor ions were isolated with a mass isolation window of 1.2 Th and fragmented by HCD at 28 % normalized collision energy. Fragment scans were acquired at 30,000 resolution with an AGC target of 5E4 and a maximum injection time of 100 ms. Selected precursor masses were excluded for the following 27 seconds.

### Protein identification

Raw mass spectral data files were searched using MaxQuant software (V1.5.7.4, (64)) using SILAC (Lys6) quantitation. Fixed modification was carbamidomethylation of cysteine; and variable modifications were oxidized methionine, deamidation of asparagine and glutamine, acetylation at protein N-terminal, and hydroxylation of proline. The data was searched against the mouse canonical Uniprot database (29/07/2015). FDR on peptide and protein level were set to 1 %. Second peptide and “match between runs” options were enabled, all other parameters were left at default settings. The mass spectrometry proteomics data have been deposited to the ProteomeXchange Consortium via the PRIDE (65) partner repository with the dataset identifier PXD023180. Reviewer account details are as follows: Username, Password: iSH0ppKX

### Statistical analysis

After MaxQuant analysis, Excel version 1.5. 21.1, GraphPad Prism 7 (GraphPad Software Inc., San Diego, CA, USA), STRING, Perseus software version 1.6.1.1 (66) and Python libraries from the Clinical Knowledge Graph’s analytics core (67) were used for data visualisation and statistical analysis.

For those proteins with a heavy to light ratio (H/L), the percentage of label incorporation was estimated as follows: (H/L/((H/L)+1)) *100. To discern protein turnover profiles in tissues, we plotted the percentage of label present in groups A versus B. In those cases were H/L ratios were not reported, we used iBAQ light (iBAQ L) and heavy (iBAQ H) data to impute H/L ratios. Missing iBAQ light in >2 samples in group A was assumed to be a consequence of fully labelled protein while missing iBAQ heavy in group B>2 samples as a result of complete turnover. H/L ratios from samples with missing iBAQ L values were imputed with the maximum H/L value + 0.1 in that column and those with missing iBAQ H values with the minimum H/L value − 0.01. To define what proteins were differentially incorporated when comparing groups D and C (unpaired t-test), we selected proteins with at least 2 valid values for H/L, and imputed missing values using the K-nearest neighbour algorithm (KNN). Further, to account for full labelling or absence of new protein incorporation, we identified proteins where H/L ratios were reported in one group but not in any of the 4 replicates in the other group. In those cases, we used heavy or light iBAQ values present in at least 2 replicates and imputed H/L ratio using uniform random values near the maximum ((maximum, maximum+std)) or the minimum ((0, minimum)) of the dataset, respectively.

The recovered proteins that showed changes in incorporation between groups were added to the STRING networks for further analyses.

The STRING networks were built using high confidence interaction score=0.7, and networks clustered to MCL inflation parameter=2 (68).

## General

We thank H. Liao and N. Ternette for useful discussions and advice with proteomics data analysis. We also thank G. Wilson for helping with figure and table preparations and revisions.

## Author contributions

Y.A-M, R. F. and T. V. were involved in the conception, design and interpretation of the data. Y. A-M conducted the animal experimental work. S. D. and Y.A-M performed sample preparation for MS. S. D. conducted MS work. A.S. D. developed code and performed data analysis. A.W., P. Ch. and R. T. participated in interpretation of the data. Y.A-M drafted the manuscript. All authors critically reviewed the manuscript and accepted final version for submission.

## Data and materials availability

All data needed to evaluate the conclusions in the paper are present in the paper and/or the Supplementary Materials. Additional data related to this paper may be requested from the authors.

