## Supplementary material for "Age-dependent changes in protein incorporation into collagen-rich tissues of mice by in vivo pulsed SILAC labelling"

**Supplementary Materials.**

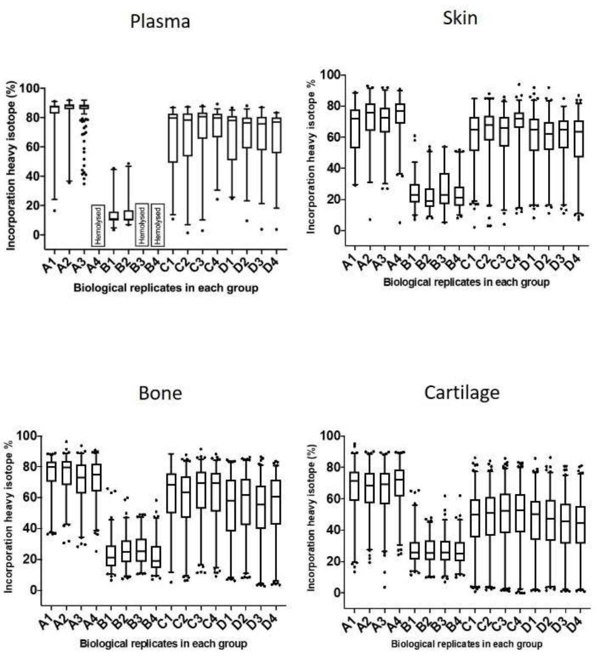


**Fig. S1**. **Incorporation of heavy isotope in plasma, skin, bone and cartilage.** Data set of individual biological replicates for all 4 groups A-D, (n=2-4). The box extends from the 25th to 75th percentiles. The line in the middle of the box is plotted at the median, and the whiskers represent the 1 and 99 percentiles.

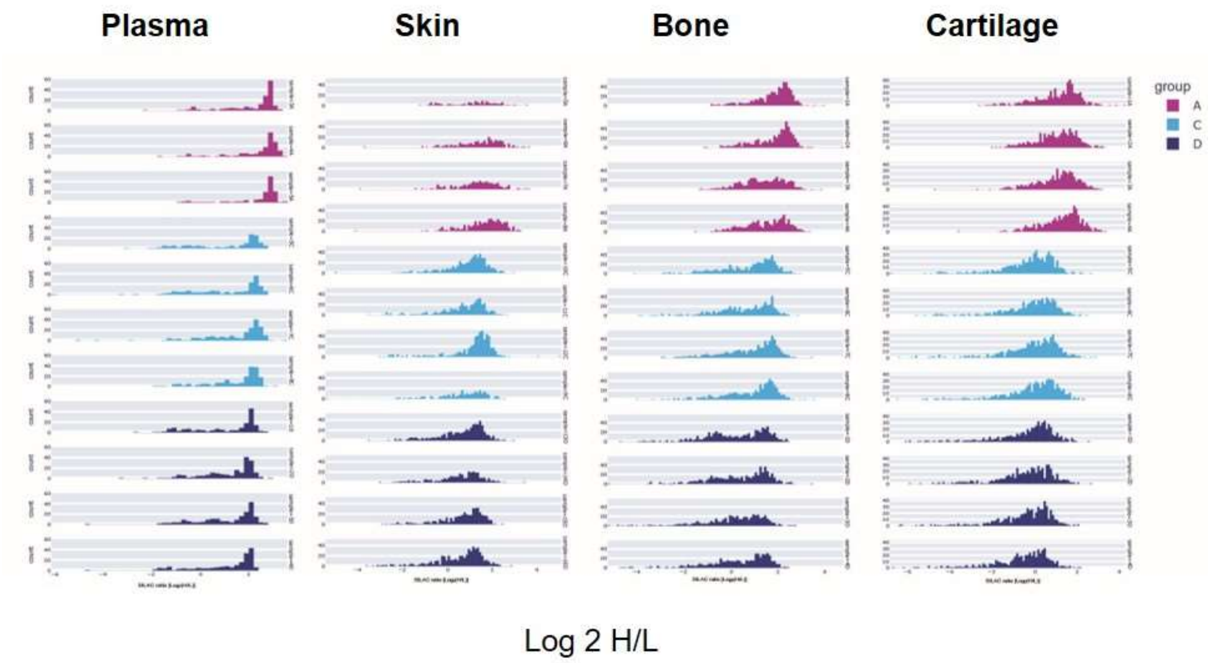


**Fig S2. SILAC incorporation into newly synthesised proteins in plasma, ventral skin, tibial trabecular bone and knee articular cartilage across the lifetime.** Log 2 H/L ratios have been plotted in the x axis. New protein synthesis was statistical significantly different between the four tissues and at all age groups, n=3-4, Kruskal-Wallis test, p<0.0001 for all comparisons.

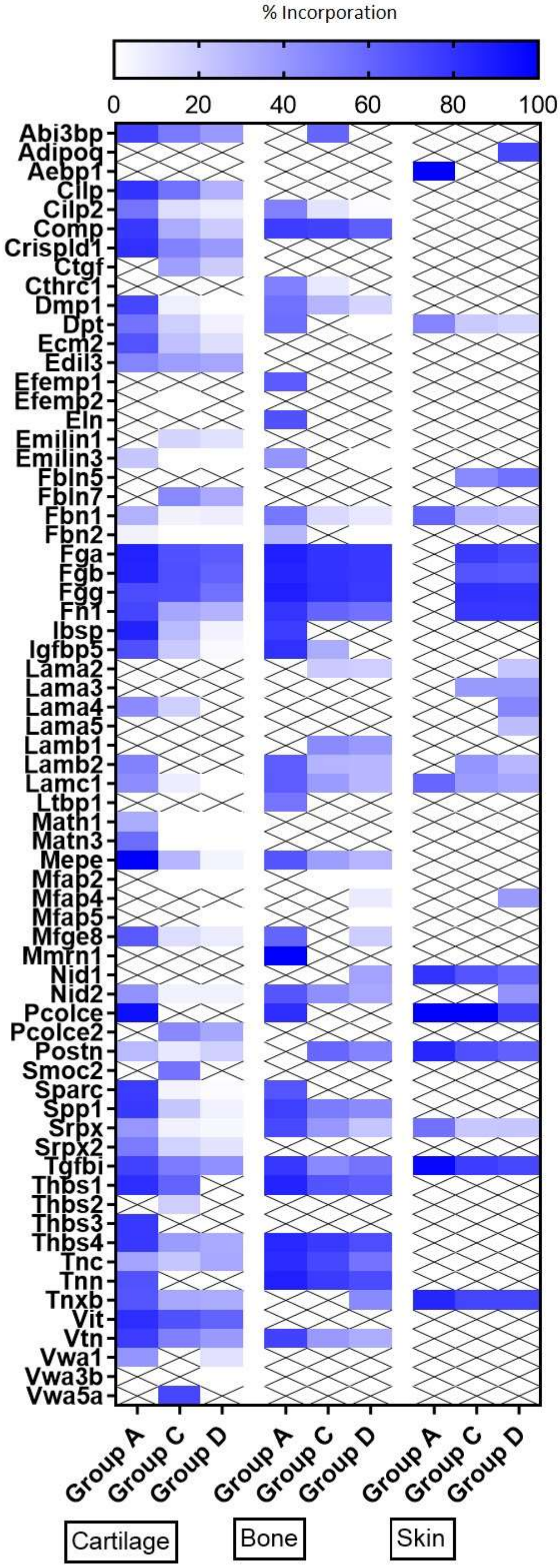
% Incorporation
0
20
40
60
80
100
Abi3bp
Adipoq
Aebp1
Cilp
Cilp2
Comp
Crispld1
Ctgf
Cthrc1
Dmp1
Dpt
Ecm2
Edil3
Efemp1
Efemp2
Eln
Emilin1
Emilin3
Fbln5
Fbln7
Fbn1
Fbn2
Fga
Fgb
Fgg
Fn1
Ibsp
Igfbp5
Lama2
Lama3
Lama4
Lama5
Lamb1
Lamb2
Lamc1
Ltbp1
Matn1
Matn3
Mepe
Mfap2
Mfap4
Mfap5
Mfge8
Mmrn1
Nid1
Nid2
Pcolce
Pcolce2
Postn
Smoc2
Sparc
Spp1
Srpx
Srpx2
Tgfbi
Thbs1
Thbs2
Thbs3
Thbs4
Tnc
Tnn
Tnxb
Vit
Vtn
Vwa1
Vwa3b
Vwa5a
Group A
Group C
Group D
Group A
Group C
Group D
Group A
Group C
Group D
Cartilage
Bone
Skin

**Fig S3. Heatmap of new glycoproteins incorporation rates into different tissues during aging.** The percentage of newly synthesized glycoproteins incorporated into articular cartilage, tibial bone and ventral skin is estimated by the percentage of incorporation of the heavy isotope ($^{13}C_6$-Lys) into the proteins during the three weeks of heavy diet. Protein synthesis and incorporation was estimated across life, covering skeletal growth (4-7 weeks old), young adults just after skeletal maturity (12-15 weeks old) and older adults (42-45 weeks old).

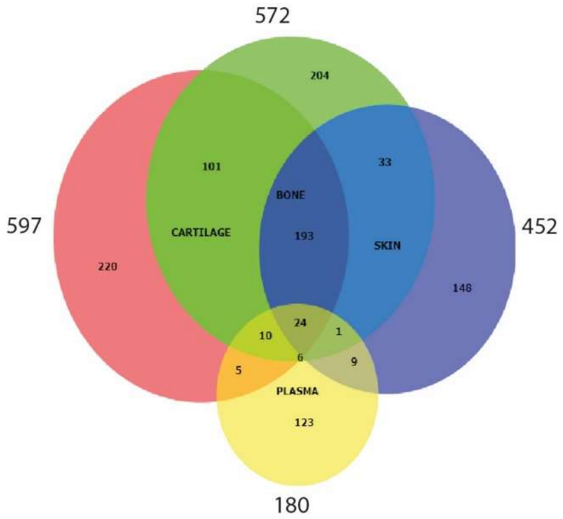


**Fig S4. Protein overlap between the four tissues.** Values represent total number of protein groups quantified in each tissue, including those proteins recovered using iBAQ light and heavy data.

**Table S1. Data supporting Fig 2. Full profile of incorporation rates (H/L), iBAQ L and iBAQ H values for plasma, skin, bone and cartilage protein groups during skeletal growth**. Groups A and B shown in Fig. 2. Values in red were imputed as follows, missing iBAQ light in >2 samples in group A, was assumed to be a consequence of fully labelled protein while missing iBAQ heavy in group B>2 samples as a result of complete turnover. Missing iBAQ light was imputed with the maximum H/L value + 0.1 in that column and missing iBAQ heavy with the minimum H/L value -0.01.

**Table S2. Data supporting Fig 3. Percentage of new collagen incorporation in the three collagenous tissues.** Cartilage, bone and skin of skeletally growing mice (group A), young adults (group C) and older adults (group D). Mean±SD, n=4, one-way ANOVA with post-Tukey's HSD test, $p<0.05$ significant.

**Table S3. Data supporting Fig 4. Percentage of new proteoglycan incorporation in the three collagenous tissues.** Cartilage, bone and skin of skeletally growing mice (group A), young adults (group C) and older adults (group D). Mean±SD, n=4, one-way ANOVA with post-Tukey's HSD test, $p<0.05$ significant.

**Table S4. Data supporting Fig S3. Percentage of new glycoprotein incorporation in the three collagenous tissues.** Cartilage, bone and skin of skeletally growing mice (group A), young adults (group C) and older adults (group D). Mean±SD, n=4, one-way ANOVA with post-Tukey's HSD test, $p<0.05$ significant

**Table S5A-D. Data supporting Fig 5 A-D. New protein incorporation across whole tissue proteomes with age.** Unpaired ttest statistics with BH corrected p values ($p<0.05$ significant) of all protein groups included in the volcano plots, plus list of proteins that were recovered using iBAQ heavy or light data.

**Table S6A-D. Data supporting Fig 5 E-H. STRING clusters with protein description.** Proteins with statistical significant changes between groups C and D (unpaired ttest BH corrected

$p>0.05$) were clustered using STRING software. Networks were built with edges high confidence interaction score of 0.7 and clustered applying MCL algorithm with inflation parameter =2.
